# On the inseparability of the prior and neural resources in behavioural bias

**DOI:** 10.64898/2026.04.07.714659

**Authors:** Henry Beale, William Harrison

## Abstract

Sensory decision making depends on environmental expectations and finite neural resources, yet these quantities are often modelled as separable values in biological systems. Here we show that this separation is neither computationally nor biologically necessary for efficient population codes. We demonstrate that prior expectations can be embedded directly in an encoding population’s tuning, thereby producing behavioural biases via Bayes-optimal inference with a uniform explicit prior. Re-analysis of V1 physiology provides preliminary support for the predicted organisation of population activity. These results highlight how systematic perceptual bias can stem from the implementation of decoding in an efficiently organised sensory code.

## Background

A long-standing goal in visual neuroscience is to understand how environmental expectations and neural resources jointly shape how the brain infers the causes of sensory input to guide cognition and action. At a mathematical level, Bayesian models specify how likelihoods and priors should combine to minimise expected loss ^1–3^. At the biological level, these quantities must be implemented through concrete mechanisms, such as neural tuning and metabolic constraints ^4–6^. Psychophysics ^7^ and neurophysiology ^8^ provide empirical constraints on how these mechanisms behave in practice. A complete account of perceptual inference must therefore link ideal computations to their neural implementation and to the biases observed in behaviour, without assuming that computational ingredients such as priors and resources remain separable across levels of explanation.

Recent work by Hahn and Wei ^9^ is a striking example of the success of information theoretic models of perceptual estimation. They formalised behavioural biases (i.e. systematic perceptual report errors) as the sum of two distinct influences: an attraction of perceptual reports toward prior expectations, and a repulsion from peaks in encoding resources. Encoding resources are derived from the Fisher information of the putative sensory encoder (see also ref ^10^). When applied across several visual estimation tasks, however, their model consistently recovered *flat* behavioural priors, with behavioural biases attributed to inhomogeneities in the resource term alone. Hahn and Wei emphasise that this outcome is mathematically permitted by their model: the loss function used in decoding perceptual estimates plays a key role in determining how the estimator trades off precision and robustness ^1^. This mathematical flexibility raises a deeper question about the biological implementation of such a trade off in sensory cortex.

Efficient coding theories propose that sensory populations of neurons internalise expectations through their tuning architecture. Prior expectations could be coded in non-uniform neural tuning density and heterogeneous widths ^1,2^, gain modulations ^5,11^, and homeostatic mechanisms that equalise metabolic load ^12^. Under a specific set of neural conditions, therefore, a sensory population of neurons may embody prior expectations in such a way that a decoding estimator need not apply an additional explicit prior ^13^. This raises a fundamental challenge for the neural implications of “flat” behavioural priors: they may not indicate the absence of expectations, but the success of an encoding strategy that has already internalised them.

Here, we formalise this argument using an efficient population code designed for visual orientation estimation. The code follows the same resource-allocation principles central to Hahn and Wei’s model, but we introduce an additional biologically motivated constraint. Simoncelli ^14^ showed that, for a population of spiking neurons, prior expectations could be embedded directly into the neurons’ encoding functions to facilitate efficient Bayesian *decoding*. The requirement for embedding is relatively trivial: the sum of the populations’ tuning curves must be proportional to the log prior over the stimulus domain. Such functional architecture would be metabolically efficient because the nervous system would not need to redundantly store the prior separately from its encoder. The resulting population simultaneously satisfies efficient encoding and efficient decoding constraints, yielding a system in which Bayes-optimal estimates are produced with a uniform explicit decoding prior.

### Sensory populations with embedded expectations

Our results, below, can be understood intuitively by considering how a biological system could implement an algorithm analogous to Bayes theorem, which is:

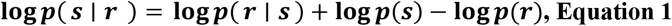

where *s* is the stimulus, *r* the neural response, *p*(*r* ∣ *s*) a property of the encoder, and *p*(*s*) the prior. Moving forward, we omit the normalising constant, *p*(*r*), without impacting the sensory estimate per se; the relative posterior depends only on the sum of the log-likelihood (log *p*(*r* ∣ *s*)) and log-prior (log *p*(*s*)).

In a biological system, the coding of prior and likelihood terms require some level of metabolic investment ^4^. One way metabolic efficiency can be achieved in a sensory system is by embedding the prior in the sensory population of neurons by adjusting the population gain to match the prior ^13,14^. Mathematically, such a biological algorithm would produce a posterior proportional to that given by Bayes theorem, but directly from the sensory likelihood. This is the case when the properties of the encoder (i.e. tuning density, width, and/or gain) are tuned so that the sum of the tuning curves, γ, is proportional to the log prior, up to an additive constant. Specifically, when:

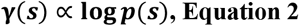

In the Supplemental Information (*An efficient population code with efficient decoding properties*), we show how and why this is the case for a population of neurons with independent Poisson noise. If we assume there is a set of encoding properties, Θ, that satisfies the above condition, then the biological likelihood generated by the encoder,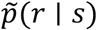, is:

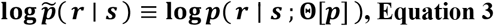

Θ[*p*] expresses that the population parameters depend on the sum of tuning curves being proportional to the log prior. We propose Equation 2 and Equation 3 as a constraint on the functional architecture of an efficient sensory encoder because it facilitates efficient decoding. For independent Poisson (or more generally log-link exponential-family) populations satisfying Equation 2, this parameterisation is equivalent to 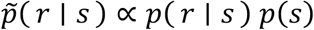: the effective likelihood naturally combines sensory codes according to expected frequencies across the stimulus domain. The relative posterior, log *p*(*s* ∣ *r*), is given directly by the effective likelihood in Equation 3 and a modified decoder with a flat (omitted) prior:

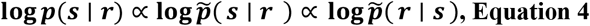

### Perceptual estimates can be reduced to a resource term with embedded expectations

We integrate the logic described above with the theory proposed by Hahn and Wei ^9^. They separate behavioural bias into two additive components. Let the true stimulus be *s*, the encoder *F*(*s*), and its Fisher information 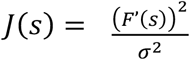, where *σ*^2^ is noise variance. Let the explicit decoding prior be *q*(*s*). For a broad class of Bayesian decoders, Hahn and Wei show that the first-order bias is:

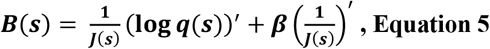

with *β* > 0 determined by the loss function. The first term of Equation 5 represents attraction to the prior, pulling estimates toward the distribution *q*(*s*). The second term reflects repulsion from the peaks of the Fisher information profile, such that non-uniformities in the allocation of resources create systematic biases in perceptual reports (see also ref ^1^). Efficient resource allocation is achieved by defining *F*(·) as the cumulative of the prior density ^5^, which has the effect of warping tuning curves so they are skewed away from peaks in the prior density (Figure 1a). This ensures that encoding resource allocation is proportional to the prior ^1,5^, thereby maximising mutual information between an environmental variable and the sensory system. In our model of visual orientation, the prior is a bimodal distribution defined as *p*(*s*) ∝ 2 − |*sin*(*s*)| (Figure 1c) ^1,2,13,15^.

**Figure 1.**
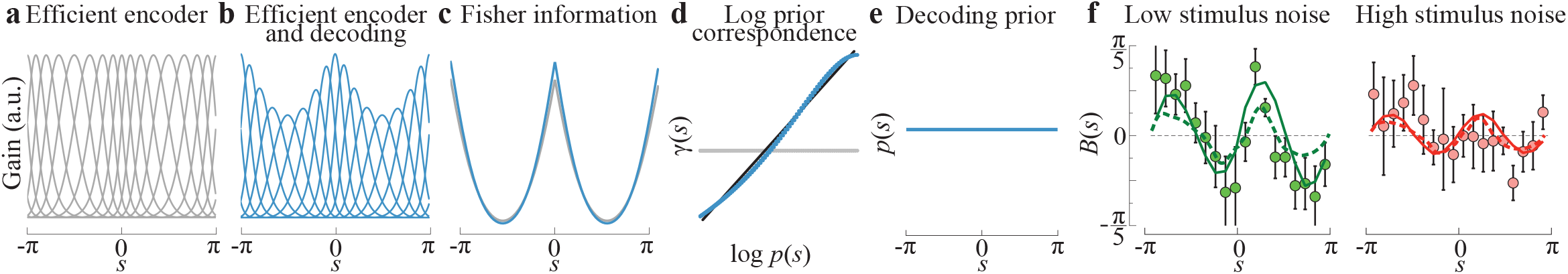
Population code design for efficient decoding with a flat explicit prior. **a)** Efficient population code in which tuning curves are warped according to the density of the environmental prior, embedding the prior in their *shape*. **b)** Efficient population code with homeostatic scaling that normalises each neuron’s total activity across the stimulus domain, embedding the prior additionally in the *gain* of each tuning curve. **c)** Both encoders allocate Fisher information proportionally to the prior, ensuring efficient resource use. Colours as per **a** and **b. d)** Grey and blue data show the sum of tuning curves for each code depicted in **a** and **b**, respectively. Note that only the normalised encoder (blue) satisfies the efficient *decoding* constraint, with the sum of tuning curves nearly perfectly proportional to the log prior (black line; shared variance = 99%). **e)** Because of this equivalence, Bayesian *decoding* is achieved with a uniform (omitted) prior, reducing Hahn & Wei’s two-term model to a single resource-dependent term. **f)** Predicted behavioural biases from the efficient decoding encoder (solid lines) capture the transition from repulsion at low noise to attraction at high noise, closely matching empirical orientation estimation data (points) and Hahn & Wei’s fits (dashed lines). Data from ^**17**^.

We designed an efficient population code of visual orientation using the same principles as Hahn and Wei. Our code is schematised in Figure 1b. Tuning curve shape is determined by the bimodal prior, but we further constrain the sum of tuning curves to be proportional to the log prior (i.e. Equation 2). We are agnostic to the mechanism that imposes this constraint, but here we use a homeostatic scaling rule ^16^, which enforces each unit to maintain a constant total spike budget under a uniform input. This rule is similar to normalisation within each unit ^5^ (i.e. equal integrated activity across the stimulus domain; see *Supplemental Information* - *Population code for visual orientation under different stimulus noise levels*). Such normalisation scales the gain of tuning curves while retaining the proportionality of Fisher information with the prior, thereby ensuring the resource is structured as per Hahn and Wei (Figure 1c). Critically, normalisation also makes the sum of the tuning curves almost perfectly correlated with the *log* prior (Figure 1d; shared variance > 99%), thereby satisfying Equation 2. This code, therefore, embeds the prior in both the shape *and* gain of the tuning curves (see also ref ^5^). Under this coding scheme, metabolic energy need not be wasted on explicitly coding prior expectations separately from the sensory encoder.

How is Hahn and Wei’s bias decomposition modified when the encoder adopts the efficient decoding properties in Figure 1b-d? First, let the population’s Fisher information be expressed as 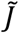 to indicate that the population resource is constrained as per Equation 2 (see also *Supplemental Information* - *An efficient population code with efficient decoding properties*). In this case, for a Bayesian decoder, the explicit *decoding* prior must be uniform (*q*(*s*) = *const*) with a derivative of 0 for all changes in *s* (Figure 1e). Equation 5 can then be reduced to a single term:

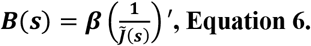

Equation 6 expresses behavioural bias solely in terms of an encoder with an embedded prior in the tuning curve shape, which determines Fisher information, and gain, which facilitates decoding. In this formulation, the loss-function exponent continues to influence the scaling captured by *β*, but it is no longer required to be tuned to generate the transition from repulsive to attractive perceptual estimate biases. That transition arises from how stimulus noise interacts with the embedded prior in the effective likelihood, rather than from adjusting the curvature of the loss itself. Thus, attraction to the prior is achieved without an explicit attraction term, and the primary degrees of freedom shift from the decoder’s loss exponent to the structure of the population code. This result is not a contradiction of Hahn and Wei’s model (i.e. Equation 5) or broader theory. Instead, Equation 6 is a special case that conforms to our hypothesised coding scheme.

We evaluated this coding scheme by fitting it to perceptual estimates from one of the visual tasks analysed by Hahn and Wei, in which observers reported the mean orientation of Gabor arrays under low and high stimulus noise^17^. For each stimulus, we generated population responses by sampling spikes from independent Poisson neurons with tuning curves defined by the double-embedded-prior architecture in Figure 1b. Decoding was performed with a maximum a posteriori (MAP) estimator using an explicitly flat (omitted) prior, such that prior expectations originated from the encoder rather than the decoder. As shown by the solid lines in Figure 1f, this model produces strong repulsion at low stimulus noise and attraction toward the prior at high noise, closely matching both the empirical data and Hahn and Wei’s original fits (dashed lines). Full details of the population code and simulation procedure are provided in the Supplemental Information.

### Neural predictions

The encoding principles schematised in Figure 1 led us to make the following predictions about the activity of real neural populations in early visual cortex. A critical function of an efficient encoder like that shown in Figure 1a is to whiten the statistics of sensory signals ^5,18^. Under this scheme, neurons should have a consistent spike rate regardless of their preferred orientations. Under the efficient decoding scheme in Figure 1b, however, the aggregate activity of a population should be heterogeneous, with higher spike rates for neurons with preferred orientations corresponding to denser regions of the prior. A comparison of spike rates across cells and sub-populations of cells tuned to different orientations can therefore provide clues about how these theoretical constraints are instantiated in biological systems.

To test the (in)homogeneity of sensory neurons’ spike counts as a function of their tuning, we reanalysed data from a meta-analysis of orientation-tuned neurons in cat V1 ^19^ (see also *Supplemental Information*). In Figure 2, we plot: a) the mean spike rates of neurons according to their preferred orientation, b) the number of cells tuned to each orientation, and c) the total mean spike rates of sub-populations of neurons with the same tuning preference. Individual spike rates are indeed whitened. However, due to the anisotropy in the number of cells tuned to each orientation, population-level activity is heterogeneous such that a decoder that aggregates spikes over a population could, in principle, employ a flat explicit prior. Whereas our model implements the efficient decoding constraint via stimulus-specific variations in gain (Figure 1b), the results in Figure 2 suggest that neural gain is constant across neurons, with prior expectations instantiated via proportionality between environmental distributions and the number of cells tuned to each orientation, at least for cat V1 cells.

**Figure 2.**
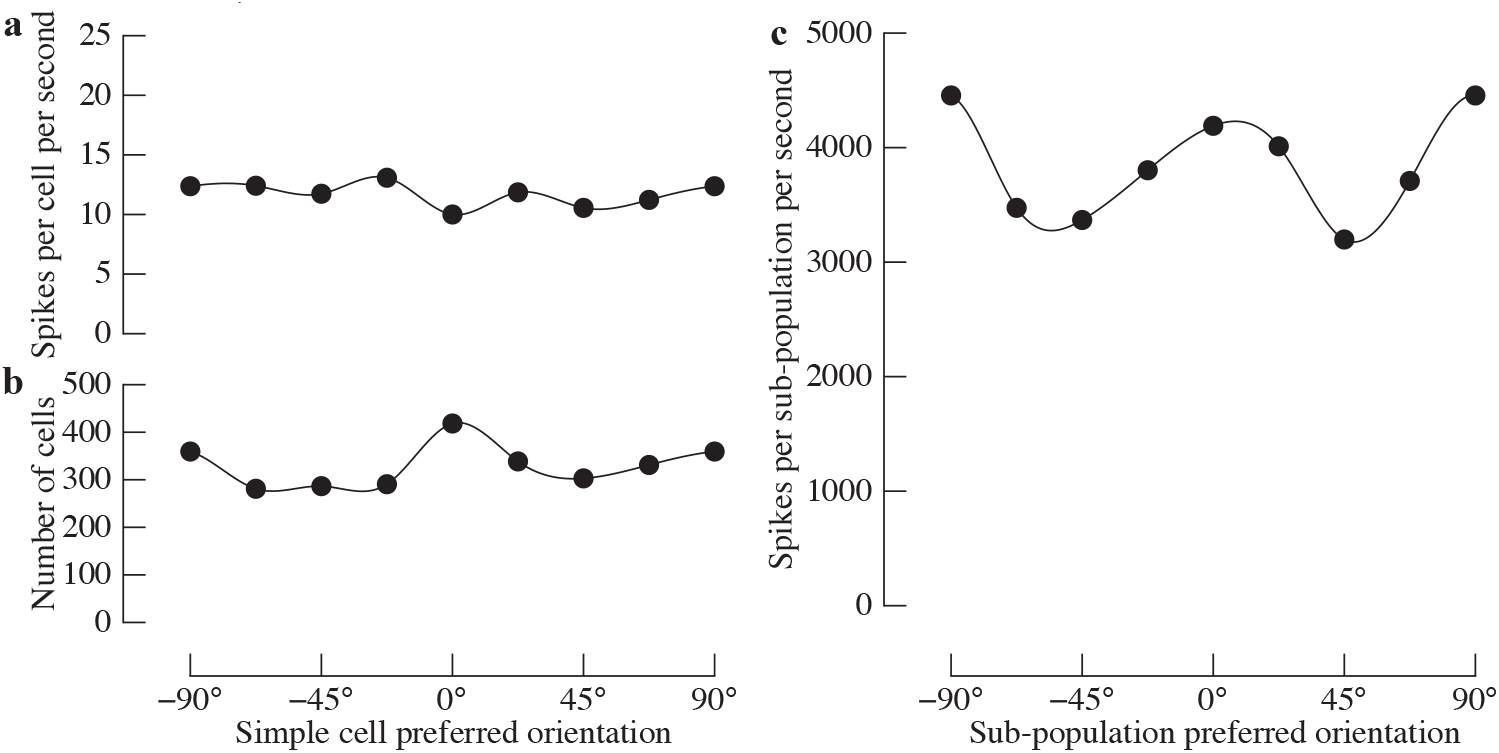
Single neuron and sub-population responses within sensory cortex as a function of orientation preference. **a)** The number of spikes per V1 neuron as a function of each neuron’s preferred orientation. **b)** The number of neurons tuned to each orientation. **c)** The cumulative number of spikes per sub-population of neurons tuned to the same orientation. Sub-population counts were computed by multiplying the mean spike rates (a) with the number of cells tuned to each orientation (b). Data taken from ^**19**^.

## Conclusion

Our modelling shows how behavioural biases can arise when a sensory population’s tuning architecture reflects environmental structure in a way that supports both encoding and decoding. By constraining a sensory population code with efficient decoding properties, a code can reproduce human biases without requiring an explicit prior that is encoded separately from the sensory likelihood. This approach offers a simple way to connect Bayesian inference to concrete neural mechanisms and naturally leads to predictions about population-level organisation in V1. Our re-analysis of existing physiology provides preliminary support for these predictions, suggesting that considering neural implementation can help clarify how perceptual biases emerge from the algorithmic structure of the sensory code itself.

## Acknowledgements

We are grateful to Dr Ivan Tomic who provided feedback on an earlier draft.

## Author Contributions

### Henry Beale

Conceptualization; Methodology; Writing - Review & Editing

### William Harrison

Conceptualization; Methodology; Formal analysis; Investigation; Writing - Original Draft; Visualization

## Supplemental Information

### An efficient population code with efficient decoding properties

Let there be a population of size *N* spiking neurons with independent Poisson noise. The population’s expected response is:

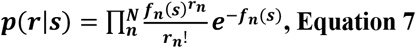

Where *f*_*n*_ is the tuning curve of the n-th neuron across the domain of ***s***. This can be expressed as log probabilities:

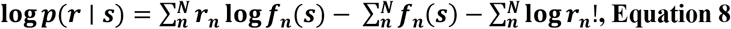

And then integrated as the likelihood in Bayes’ theorem (Equation 1):

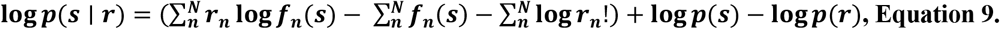

The parenthetical term in Equation 9 is the Poisson likelihood (i.e. Equation 8), while the remaining terms are the prior and normalising constant. We can simplify by removing any terms that do not depend on the stimulus:

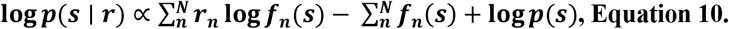

From Equation 10, it is clear that the second and third terms, which correspond to the sum of the tuning curves and the log prior, respectively, would cancel out when:

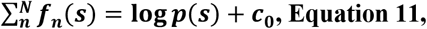

which restates Equation 2. When this constraint is met, the posterior for a given neural response is given by a modified decoder with flat (omitted) prior:

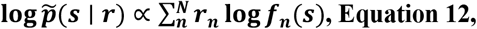

up to an additive constant. To emphasise the simplicity of this coding scheme, the posterior is proportional to the effective sensory likelihood with an embedded prior:

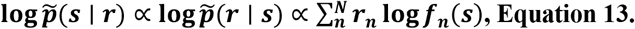

### Population code for visual orientation under different stimulus noise levels

The population code shown in Figure 1a includes 64 idealised units with circular Gaussian tuning functions. Changing the number of units or sampling resolution does not change the results. Following Wei and Stocker ^1^, we assume that these tuning curves are symmetric and evenly spaced in sensory coordinates, and we set the tuning widths to be equal across the population (kappa = 7). When transformed into stimulus coordinates, the shapes of the tuning curves are re-allocated according to the cumulative density of the prior, which we specified as:

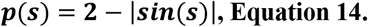

Homeostatic scaling normalises each tuning curve to integrate to the same value. This was achieved by:

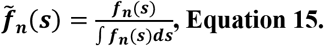

Fisher information for the population code is:

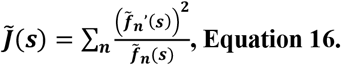

To generate model predictions, we discretised orientation space into 360 points on [−π, π) and simulated a population of 64 units with equally spaced preferred orientations in sensory coordinates. For each orientation condition, we modelled the mean stimulus as a von Mises distribution centred on the true orientation *s* with concentration κ_*ext*_, corresponding to the low-noise and high-noise conditions in Tomassini et al. ^17^ (2° and 14° orientation SD, converted to κ_*ext*_ using the CircStat toolbox ^20^). The model predictions for each observer were determined by two free parameters that influenced the mapping between the encoded population response and behavioural estimates. The first parameter was a global gain factor, ***g***, which scaled the expected spike rates of all neurons and therefore controlled the signal-to-noise ratio of the population response. The second parameter was an external-noise scaling factor, ***α***, which multiplied the concentration of the von Mises target distribution used to generate population activity. Given a target distribution *p*_*target*_(*s*) and tuning curve ***f***_***n***_(*s*) for neuron ***n***, expected spike counts were computed as:

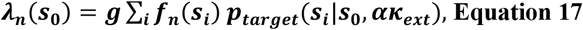

where *s*_*i*_ denotes the discretised orientation value at position *i* on the grid. As described below, a single pair of parameter values (***g, α***) was estimated per participant and applied to both the low- and high-noise conditions. Therefore, the free parameters could not differentially shape bias in one noise level vs the other, ensuring that the model’s noise-dependent changes in bias arose from the encoder and not from parameter variation. For each combination of target orientation ***s***_**0**_, external-noise concentration **κ**_***ext***_, gain ***g***, and noise-scaling factor ***α***, spikes were generated as independent Poisson samples:

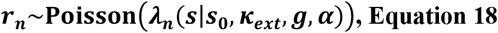

with an optional baseline rate (set to zero in the reported fits). Each response vector was decoded on the discretised orientation grid using a maximum a posteriori (MAP) estimator with a flat explicit prior. The log-likelihood of orientation ***s*** given response ***r*** is:

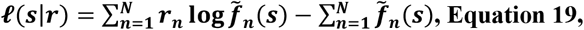

up to an additive constant that does not depend on ***s***. Note that 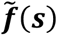 defines the scaled tuning curves (see Equation 15). In general, the log-posterior combines the log-likelihood, log prior, and constant:

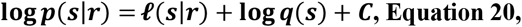

where *q*(*s*) is the decoder’s explicit prior and *C* is a normalising constant. By design, however, the decoder prior for our model is uniform (*q*(*s*) = const), so log *q*(*s*) is absorbed into C, and the MAP estimate reduces to a pure likelihood maximisation:

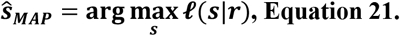

Thus, the decoding stage omits any explicit prior term, and all prior structure influencing the estimate arises from the embedded prior in the population code. For each condition, the model’s predicted estimate was obtained as the circular mean of the 2,000 MAP estimates, and bias was computed as the circular difference between this mean and the true orientation. For each parameter pair, we generated model biases for all stimulus orientations and both noise levels, and compared these values to the empirical bias functions for each observer. The goodness of fit was quantified as the sum of squared circular deviations across orientations and stimulus noise conditions. The best-fitting gain and noise-scaling parameters for each participant were defined as those that minimised this error, and these values were used to generate the predictions displayed in Figure 1f.

### Analysis of cat V1 activity

We re-analysed data from a meta-analysis of cat V1 responses, by Li et al. ^19^. We used WebPlotDigitizer ^21^ to extract the reported activity of V1 simple cells tuned to high spatial frequencies, which showed the strongest cardinal biases. Li et al. reported the directional tuning of these neurons in response to drifting gratings, but also found that the majority were circularly symmetric (i.e. a neuron tuned to 0° direction also responded to 180° direction). We therefore averaged activity across neurons tuned to polar opposite directions. Figure 2 simply shows the mean activity, the number of neurons, and these values multiplied to show the total spikes across the population.

## Footnotes

1 Key to Hanh and Wei’s model is that stimulus noise interacts with the loss exponent to reduce or eliminate the behavioural influence of an explicit prior, allowing the model to fit the data with a flat prior if and when the resource term already reflects structured environmental expectations.

